# Telomere length can be estimated from fish scales

**DOI:** 10.64898/2026.01.07.698083

**Authors:** Gaëlle Brahy, Aurélie Manicki, François Gueraud, Maika Minjou, Sylvie Oddou-Muratorio, Jacques Labonne

## Abstract

Telomeres, repetitive DNA sequences at the ends of chromosomes, are increasingly studied as biomarkers of physiological stress as well as ageing processes. In ichthyology, fish scales provide valuable information regarding life history and ageing and are thus often preserved in substantial collections. This study aims to adapt a quantitative Polymerase Chain Reaction (qPCR) method to estimate relative telomere length (RTL) from fish scales archived in such collections. Using the brown trout (*Salmo trutta*) as a model species, we show that archived scales contain enough DNA to perform qPCR-based RTL measurement. RTLs in scales are on average higher than in fins, however we found no significant correlation between RTL values of both tissues at the individual level. Analysing archived scales originating from the subantarctic Kerguelen Islands introduced brown trout populations, we find substantial variation in RTL between rivers, age classes and body size.

## INTRODUCTION

Telomeres are highly repetitive genomic regions located at the ends of chromosomes where they prevent chromosomal fusion and protect against chromosomal erosion. This protective function is crucial because the incomplete replication at the 3’ extremity of the lagging strand is responsible for telomeres shortening with each cell division cycle. Excessive telomere shortening, such as caused by ageing or stress exposure, is a key driver of cell senescence (Meyne *et al*., 1989; Blackburn, 2000; Shay and Wright, 2019).

Telomere length is used as a biomarker of age and/or stress in numerous taxa (Monaghan, 2014; Spurgin *et al*., 2018; Burraco *et al*., 2022, McLennan *et al*., 2022). Telomere shortening with age, associated with telomere erosion at each division cycle, is widely acknowledged in mammals and birds (Haussmann and Marchetto, 2010; Spurgin *et al*., 2018; Demanelis *et al*., 2020). In ectotherms however, and in fishes in particular, results are more contrasted: while some studies report an ageing effect (Hatakeyama *et al*., 2008; Hartmann *et al*., 2009; Evans *et al*., 2021), other find no trend (Lund *et al.,* 2009; Pauliny *et al.,* 2015) or even variable trends within a same species (Simide *et al.,* 2016). These differences between and within studies could either be due to differential telomerase activity between tissues, life stages and taxa (Hatakeyama *et al*., 2008; Hartmann *et al*., 2009) and/or life stages (Lund *et al*., 2009; Smith *et al*., 2022) or to oxidative stress (von Zglinicki, 2002, Haussmann *et al.,* 2003; Monaghan, 2010). In short, telomere length variation is triggered by a number of factors and processes, many of them being of interest for biodiversity dynamics and conservation.

Relative telomere length (RTL) can be routinely estimated using real-time quantitative Polymerase Chain Reaction (qPCR), a cost-effective method that allows to process large numbers of samples (Cawthon, 2002; Lai *et al*., 2018). This approach up to now required potentially crippling or lethal sampling methods. For instance, in salmonids, studies used DNA extracted from fins (Näslund *et al*., 2015; Pauliny *et al*., 2015; McLennan *et al*., 2016; McLennan *et al*., 2022; Duncan *et al*., 2023) or other tissues (cross sections containing muscle, liver, skin and blood; Debes *et al*., 2016). To our knowledge, scales were never used for RTL estimation, whereas they are a popular tissue for fish studies, notably because their removal has a low impact on fish welfare and behaviour and because they offer the lowest cost of conservation. They are thus increasingly archived in extensive banks, spanning a large spectrum of species, locations, ecosystems and dates (Chagnoux, 2024; Mougin *et al*., 2018). Scales allow the collection of many scientific data (age and growth determination, microchemistry, isotopic and molecular biology analyses). Scales samples typically contain DNA from a limited number of dermal and epidermal cells (Nielsen *et al*., 1999; Rakers *et al*., 2010). Our objective is to propose and validate a novel method to estimate RTL via qPCR, using DNA extracted from archived fish scales, with brown trout (*Salmo trutta*) as the test species. First, we adapted the existing qPCR-based protocol to this specific biological material − a technological challenge given the limited number of epidermal cells typically present in scales samples. Next, we investigated tissue-type correlation of RTL at the individual level, by comparing measurements from both scales and fins samples in a benchmarking perspective. Finally, to illustrate the potential applications of our method, we explored the variation of RTL from scales samples of brown trout individuals, spanning multiple rivers, age classes and body sizes.

## MATERIEL & METHODS

### Sampling design

*S. trutta* is a facultative anadromous species, presenting a freshwater and a seawater ecotype. To ensure the population of origin, we only focused on the freshwater ecotype, selecting individuals smaller than 200 mm and/or younger than 4 years old (age noted 1+, 2+ or 3+). For tissue-type comparison, we used archived scales and fins from 36 individuals of the Nivelle river in the South West of France, sampled between 2022 and 2025 (Suppl. Tab. S1a). To investigate RTL variation in scale samples, we used archived scales originating from 446 individuals from seven rivers of the subantarctic Kerguelen islands, sampled between 2002 and 2019 (Suppl. Tab. S1b). All samples are stored in the Colisa repository (https://colisa.fr).

### Samples preparation

To constitute biological tissue collections, fins samples are usually collected by clipping out a section of the anal fin, and stored in pure ethanol. Scales samples are generally obtained by scraping body surface with a blade, and storing this material in a paper envelope protected from light exposure and at room temperature. When drying, this material attaches to the surface of the paper, forming a stain hereafter called scales zone.

We prepared fins samples by taking out approximately a 3 mm² section of the archived fin sample for each individual. For tissue-type comparison, in order to preserve the very small amount of archived tissue available, only two scales for each individual were used. For Kerguelen scale samples, a preliminary analysis comparing three sizes of scales zones (4, 8 and 16 mm²) showed that an 8 mm² area was enough to ensure DNA quality and quantity optimization (result not shown). An area of 8 mm² contains between 100 and 50 scales, typically for a one-year-old individual. We therefore cut out an 8 mm² section of the scales zone (including the paper envelope), and placed it directly into Eppendorf tube for lysis, thereby avoiding biological material loss and contamination through manual handling. Any DNA unintentionally extracted from mucus microbial communities is not amplified by qPCR, since bacteria do not have telomeres and yeasts have different telomere sequences (Fulnečková *et al*., 2013; D’Angiolo *et al*., 2023).

### DNA extraction and quality control

DNA was extracted with the Nucleospin Tissue kit (Macherey Nagel) according to the supplier’s recommendations with an overnight lysis step. DNA samples were eluted in 28 µL of Elution Buffer to obtain the highest possible DNA concentration. Kerguelen DNA samples were purified using the OneStep PCR Inhibitor Removal kit (Zymo Research) to get rid of PCR inhibitors. DNA concentrations were quantified with 1 µL of each DNA sample using the Qubit™ ds DNA BR assay kit (Invitrogen). The quality of DNA was evaluated by electrophoresis on 1.5 % agarose gel. DNA was then diluted with ultra-pure DNase free water at 1 ng·µL^−1^ for the qPCR analyses.

### Real-Time Quantitative Polymerase Chain Reaction (qPCR)

Details on the RTL estimation through qPCR principle can be found in Pfaffl (2001) and Cawthon (2002). We amplified the DNA of telomere and β-actin, the reference gene, following the qPCR protocol of Näslund *et al*. (2015) with some modifications. First, qPCRs were performed in a volume of 5 µL with a final concentration of 0.2 ng·µL^−1^ DNA to preserve DNA collection while using DNA concentrations consistent with Näslund *et al*. (2015). Second, the telomere qPCR program was adjusted as follows: after an initial denaturation at 95 °C during 10 min, we ran 50 cycles, each including (1) 15 s at 95°C (denaturation), 30 s at 56°C (primer hybridization) and 30 s at 72°C (elongation). The β-actin qPCR program started with the same initial denaturation and followed with 45 cycles each including 15 s at 95°C (denaturation) and 1 min at 56°C (primer hybridization). Elongation took place during the transition from 56°C to 95°C. Both programs were completed with a melting curve (+ 0.5°C, 65°C to 97°C, 2.5°C·sec^−1^) to evaluate PCR specificity. We manually prepared qPCR mix using the LigthCycler 480 SYBR Green Master kit following the recommendations of the manufacturer (Roche). Deposits were realized by the same person to avoid operator bias. qPCRs were performed using a LigthCycler 480 thermocycler (Roche). qPCRs of telomere and β-actin were prepared simultaneously, loaded on separate plates and run on the same day to minimize technical bias.

Samples were deposited on five different 384-wells-plates grouped by tissue type, decade of the collection year, and individual age (Suppl. Tab. S2). Firstly, DNA of focal samples were loaded in duplicate, and calibrator samples were run in triplicate on each plate. Secondly, to evaluate qPCR efficiency on each plate, a standard curve was generated from five serial dilutions of a standard sample (1:10 dilutions, ranging from 15 to 0.0015 ng·µL^−1^); this standard scale was replicated four times.

Thirdly, to enable comparison between plates, four calibrator samples were loaded in triplicate on each plate. These calibrators were selected to provide the best possible representation of river of origin, collection year and individual age.

### RTL estimation

qPCR data results were analysed using LightCycler® 480 Software version 1.5.0 to assess amplification efficiency and error rates for each qPCR run and to extract the cycle threshold (C_t_) values for all samples. Melting curve (fusion peak) analyses were examined to exclude samples showing evidence of contamination or primer-dimer formations. C_t_ values were determined using the E-method with the Second Derivative Maximum approach.

We excluded samples for which amplification failed for at least one of the duplicates (C_t_ > 15 for telomere or C_t_ > 27 for β-actin). Additionally, we removed samples showing a C_t_ difference between duplicates greater than 0.6. The RTL of each focal sample was calculated using the formula of Cawthon (2002) adapted by Naslünd *et al*. (2015):

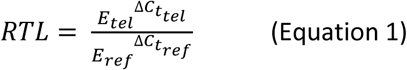

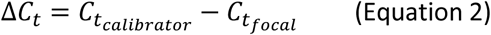

Where *E*_*tel*_ and *E*_*ref*_ are the qPCR efficiency value for telomere and β-actin respectively and 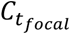 is the average of the C_t_ values of each focal sample’s duplicates. For each calibrator, the two closest C_t_ values among the triplicates were averaged, with difference between these values never exceeding 0.6. The mean C_t_ value across the four calibrator samples provided a single calibrator C_t_ value 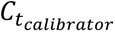 per plate. Thus, 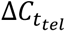 represents the telomere C_t_ value of the focal sample relative to the average telomere C_t_ value of the calibrators on the considered plate. Similarly, 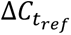 represents the β-actin C_t_ value of the focal sample relative to the average β-actin C_t_ value of the calibrators on the considered plate.

### Statistical analyses

For tissue-type comparison, we investigated difference between RTL values distributions of fins and scales using a Wilcoxon test for paired data, and explored the co-variation of RTL values between both tissues using a Spearman correlation test. To investigate the variation of RTL according to river and individual age (categorial factors), and body size (continuous, standardized), we used a generalized linear model (GLM) with a Gamma distribution and a log link. Model parameters were tested using Wald z-tests, and overall effects were assessed using likelihood ratio χ² tests based on analysis of deviance.

## RESULTS AND DISCUSSION

### DNA extraction

For Nivelle samples, DNA was successfully extracted from all 36 fins samples, and from 33 out of 36 scales samples, failure being likely due to the very small amount of tissue available. DNA concentrations obtained from fins samples, ranging from 4.4 to 220 ng·µL^−1^ with an average of 90.21 ng·µL^−1^, were higher than those obtained from scales samples, ranging from 4.08 to 27.2 ng·µL^−1^ with an average of 7.44 ng·µL^−1^ (Suppl. Tab. S1a). This difference reflects the relatively smaller amount of cell material contained in scales as compared to fins.

For Kerguelen samples, DNA was successfully extracted from 442 out of 444 scales samples, and concentrations ranged from 3.8 to 438 with an average of 67 ng·µL^−1^ (Suppl. Tab. S1b). DNA concentration varied significantly between rivers (Kruskal–Wallis test: χ² = 411.59, df = 6, p-value < 2.2 10⁻¹⁶), and individual age (Spearman’s ρ = −0.33, p-value < 2.2 10⁻¹⁶) but did not appear to be significantly related to collection year (Spearman’s ρ = −0.061, p-value = 0.063). We note that individual ages are not distributed evenly between rivers.

### qPCR amplification

Efficiency of qPCR were similar for DNA extracted from scales or fins. Error rate were also comparable across all qPCR runs targeting the same gene (telomere or β-actin) and were robust regardless of tissue type and sample age (0.005 to 0.021 for telomere; 0.005 to 0.023 for β-actin; Suppl. Tab. S2).

In the Nivelle dataset, only one sample failed to meet the C_t_-difference threshold, leaving 32 individuals (97%) with reliable C_t_ values for RTL estimation (Suppl. Tab. S3a). In the Kerguelen dataset, 16 samples failed to meet the C_t_-value threshold, and 115 failed to meet the C_t_-difference threshold, resulting in 311 individuals (71%) with reliable values for RTL estimation (Suppl. Tab. S3b). Higher C_t_ differences were observed for telomeres as compared to β-actin (Suppl. Fig. S1). The proportion of samples with C_t_ difference < 0.6 among replicates tended to increase with time since collection year in Kerguelen samples, but not significantly (Spearman’s ρ = 0.54, df=9, p-value = 0.09). In the retained samples, C_t_ values ranged from 9.47 to 14.10 (telomere) and from 21.57 to 25.68 (β-actin; Suppl. Fig. S2).

### Comparison of RTL between tissue types

RTL values were significantly lower in fins than in scales (Fig. 1), as confirmed by a Wilcoxon test for paired data (N =32, V = 84, p-value = 4.3 10^−4^, r = −0.59). These results indicate that RTL measurements from scales are of comparable magnitude to those from fins, suggesting that using qPCR to estimate RTL from scales is effective. At the individual level, no correlation between tissue types was detected (Spearman’s rho = 0.18, p-value = 0.31).

**Figure 1.**
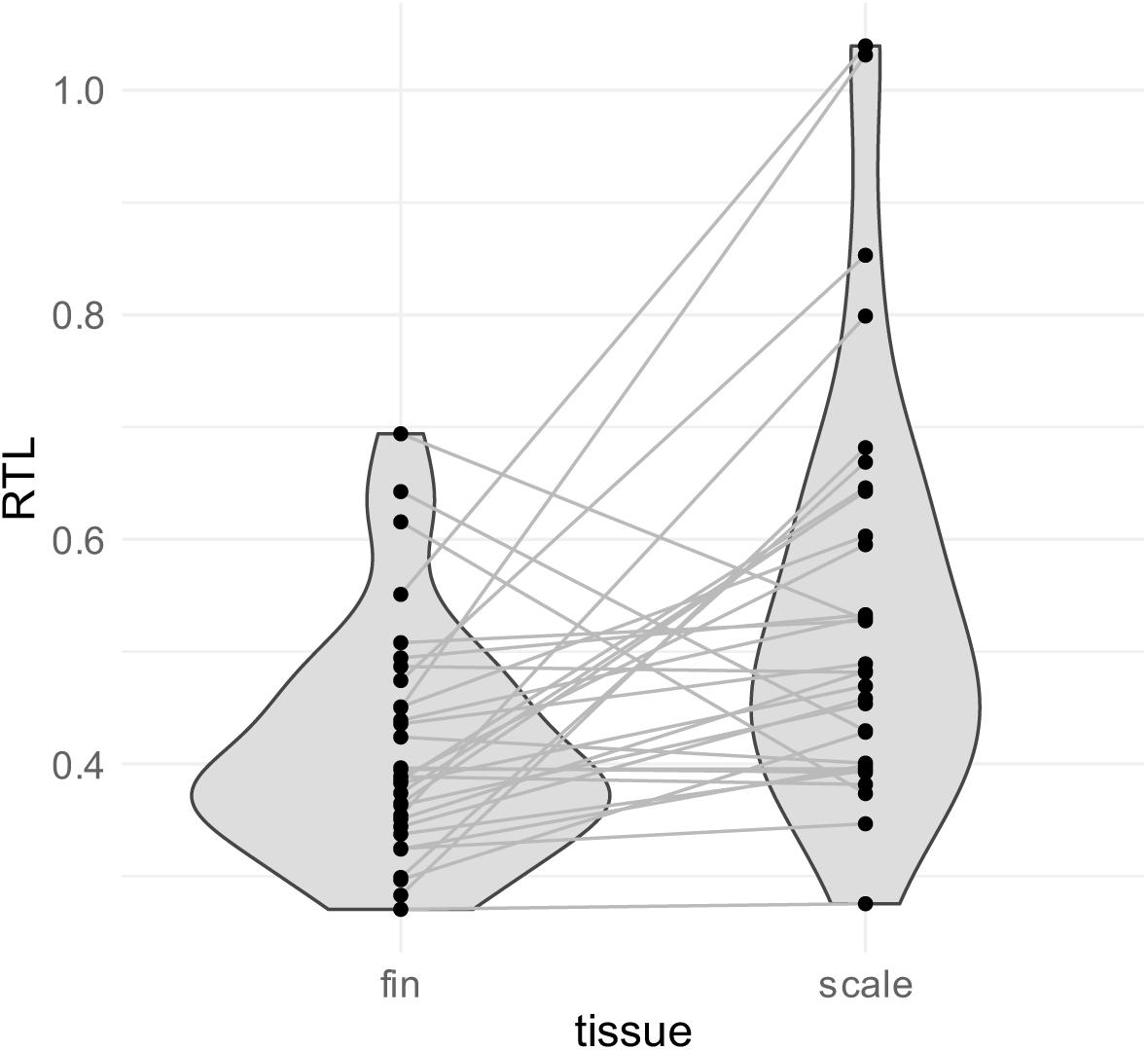
Comparison of RTL estimated from fins and scales samples in 32 brown trout individuals from the Nivelle river. Points represent RTL values obtained for each tissue type. Lines connect RTL values from the same individual, highlighting tissue dependent variation.

The observed differences between RTL estimated from different tissue likely arise from several non-exclusive factors. First, part of the differences may originate from tissue-specific storage conditions. While fins were stored in alcohol, scale samples were dried, and both tissue types were kept at room temperature. Such difference can affect DNA quality and quantity, potentially influencing qPCR performance (Lin *et al*., 2019).

Second, these differences may reflect intrinsic biological difference between tissues, including variation in telomerase activity and/or differential responses to physiological stress. Interestingly, Debes *et al*. (2016) reported tissue-dependent relationships between RTL and growth: in brown trout, among six tissue types, fins displayed the weakest RTL-growth association, whereas muscle showed the strongest. This suggests that tissue types undergo different levels of oxidative stress for a given trait value, leading to tissue-dependant correlations between RTL and traits. Similarly, Demanelis *et al*. (2020) reported that exposed skin had the second-longest telomere among twelve human tissue types (after testis). By analogy, fish epidermal tissue may exhibit higher telomerase activity, particularly as it contains undifferentiated cells that sustain renewal of the tissue (Rakers *et al*., 2010). This could explain the higher RTL values observed in scale-derived DNA. Indeed, although fin tissue also contains epidermis, it likely contains a lower proportion of epidermal cells relative to cartilaginous cells, which have a slower turnover rate. While studies in humans generally report mostly positive – though often weak – correlations between tissue types (Stout *et al*., 2017; Goldman *et al*., 2018; Demanelis *et al*., 2020), tissue-specific patterns of RTL variation in fish may reflect differences in cell composition and telomerase activity. Although this prevents direct comparison of RTL estimated from different tissues, this pattern opens new perspectives to study tissue-specific impact of oxidative stress.

### Variation in RTL among rivers, body size and individual ages

In the Kerguelen data set, we successfully measured RTL from 311 individuals, out of which two values were considered as outliers (RTL > 3) and excluded. A simple GLM model fitted on these 309 individual values (Fig. 2) explained 64% of observed variance and indicates that RTL differed significantly among rivers and age classes, with a significant interaction between these two factors (all p-values < 1 10^−6^, see Suppl. Tab. S4a). RTL were lower for older fish, and once removed the effect of age, body size had a positive effect on RTL (see parameter estimates Suppl. Tab. S4b). These snapshot results obtained from archived scales suggest that ample variation in RTL can be found in samples from natural populations, potentially correlating to variables of interest for ecologists.

**Figure 2.**
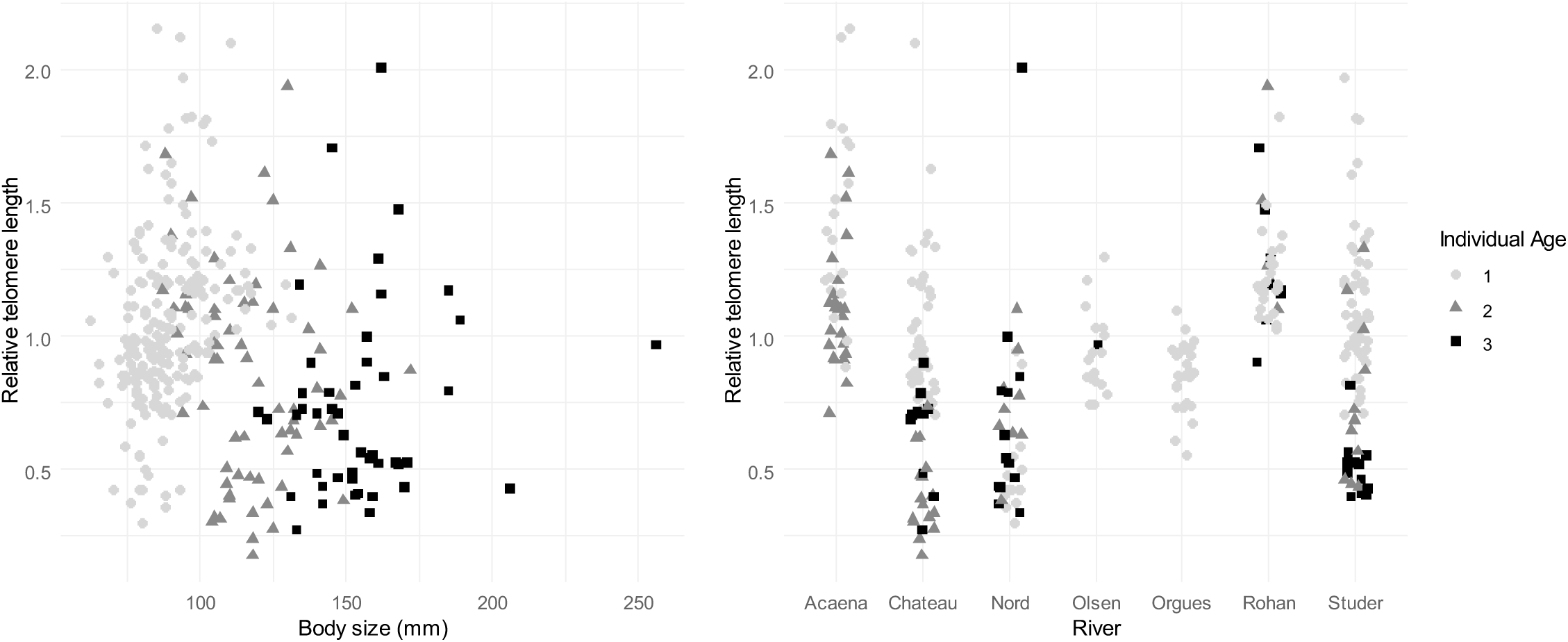
Variation of RTL values in Kerguelen scale samples, depending on body size (x-axis, left) or river of origin (x-axis, right). Individual age is represented by colours.

Of particular interest is the fact that, in fish, age can often be estimated from scales, sometimes with a good accuracy (especially in young fish in some species as is the case here, Rifflart *et al*., 2006). Moreover, size-at-age can also be estimated from scales. Combined with this knowledge, our study therefore offers an exciting avenue to analyse how growth relative to age influences RTL variation, whereas in many other non-fish species, age estimation is so challenging that RTL often serves as a proxy of age. We therefore hope that this minimally invasive, cost-effective method will open up many opportunities to analyse samples and data from widely available past and current collections, enabling to investigate questions pertaining to stress and aging processes across a broad range of fish species.

## ACKNOWLEDGEMENTS

We thank the French Polar Institute for the long-term support in the sampling effort of fish scales in the subantarctic Kerguelen Islands. We also thank F. Lange for providing fins and scales tissues from the Nivelle river. We are grateful to E. Plagnes, S. le Garrec, V. Veron and C. Houdelet for precious advice on the adjustment of qPCR protocol. The PhD of G. Brahy is funded by the EDENE program, that has received funding from the European Union’s Horizon 2020 Research and Innovation Program under the Marie Skłodowska Curie Actions-Grant agreement N° 945416. This work was also partly supported by INRAE ECODIV division (TELOSCALE project).

## DATA AVAILABILITY

Data for RTL comparison between tissues and data for RTL variation among rivers, body size and individual age are available at https://doi.org/10.57745/FX83AD

## SUPPORTING INFORMATION

Supplementary materials available in the file: Brahy et al., 2026_supp_mat.pdf

## CONTRIBUTIONS

A.M. designed the first qPCR protocol. F.G. and G.B. prepared the scales samples. M.M., A.M. and G.B. performed the biomolecular analysis. G.B., A.M., M.M., J.L. and S.O.M. analysed the data. G.B. and A. M. wrote the first draft of the manuscript and all the authors contributed to the final version.

