## Supplementary materials for "Telomere length can be estimated from fish scales"

### Supplementary Table S1. Sampling design and average DNA concentration (red italic) for a) tissue-type comparison and b) investigation of RTL variation in Kerguelen scale samples.

In Tab. S1b, DNA concentration varied significantly between rivers (Kruskal–Wallis test: $\chi^2 = 411.59$, df = 6, p-value < 2.2 $10^{-16}$), and individual age (Spearman's $\rho = -0.33$, p-value < 2.2 $10^{-16}$) but did not appear to be significantly related to collection year (Spearman's $\rho = -0.061$, p-value = 0.063). We note that individual ages are not distributed evenly between rivers.

a)

| **Tissue type** | **Nb of samples for DNA extraction** | **Nb of samples after extraction with *their average DNA concentration (ng·µl$^{-1}$)*** | **Collection year of successfully extracted samples** | | | |
|---|---|---|---|---|---|---|
| | | | **2022** | **2023** | **2024** | **2025** |
| Fins | 36 | 36<br>*90.21* | 3<br>*79.7* | 5<br>*58.16* | 21<br>*87* | 7<br>*127.2* |
| Scales | 36 | 33<br>*7.44* | 3<br>*5.78* | 5<br>*6.8* | 20<br>*6.62* | 5<br>*12.26* |

b)

| **River** | **Collection year** | **Nb of samples for DNA extraction** | **Nb of samples after extraction *with their average DNA concentration (ng·µl$^{-1}$)*** | **Nb of samples after extraction per age class, *with their average DNA concentration (ng·µl$^{-1}$)* when relevant** | | |
|---|---|---|---|---|---|---|
| | | | | **1+** | **2+** | **3+** |
| Acaena | 2018 | 35 | 35<br>*123.64* | 3<br>*108.3* | 32<br>*125* | 0 |
| | 2003 | 25 | 25<br>*73.59* | 25 | 0 | 0 |
| Chateau | 2018 | 66 | 66<br>*55.9* | 25<br>*102* | 21<br>*23.2* | 20<br>*32* |
| | 2001 | 31 | 31<br>*35.84* | 31 | 0 | 0 |
| Nord | 2010 | 60 | 58<br>*20.45* | 19<br>*17.6* | 20<br>*26.6* | 19<br>*16.75* |
| Olsen | 2018 | 25 | 25<br>*32.88* | 24<br>*30.7* | 0 | 1<br>*83.6* |
| Orgues | 2018 | 31 | 31<br>*75.89* | 31 | 0 | 0 |
| Rohan | 2019 | 32 | 32<br>*74.68* | 13<br>*91.7* | 8<br>*75.2* | 11<br>*54.1* |
| | 2002 | 35 | 35<br>*113.83* | 35 | 0 | 0 |
| Studer | 2011 | 70 | 70<br>*94.39* | 32<br>*151.95* | 19<br>*33.42* | 19<br>*58.43* |
| | 2002 | 34 | 34<br>*197.2* | 34 | 0 | 0 |
| **Total** | | **444** | **442** | **272** | **100** | **70** |

### Supplementary Table S2. Plate design, efficiency and error rate of qPCR for telomere and β-actin.

The five qPCR plates differed in terms of individual age, tissue type and collection year. Note that plate 1 includes Nivelle focal samples – for tissue-type comparison analysis, while plates 2 to 5 include Kerguelen focal samples – for investigation of RTL variation.

| n° qPCR plate | Individual age or size | Tissue | Collection year | Telomere | | β-actin | |
|---|---|---|---|---|---|---|---|
| | | | | Efficiency | Error rate | Efficiency | Error rate |
| plate 1 | < 200 mm | scales - fins | 2022-2025 | 1.769 | 0.005 | 1.929 | 0.009 |
| plate 2 | 1+ | scales | 2018-2019 | 1.787 | 0.014 | 1.78 | 0.008 |
| plate 3 | 1+ | scales | 2011-2001 | 1.794 | 0.012 | 2.04 | 0.019 |
| plate 4 | 2+ and 3+ | scales | 2018, 2010, 2011 | 1.864 | 0.011 | 1.984 | 0.005 |
| plate 5 | 2+ and 3+ | scales | 2018, 2010, 2011 | 1.857 | 0.021 | 2.052 | 0.023 |

### Supplementary Table S3. Results of the qPCR analysis showing the number of samples that passed the two quality thresholds for a) tissue-type comparison and b) investigation of RTL variation in Kerguelen scale samples.

The first quality threshold retained samples for which real time qPCR yielded below-threshold $C_t$ estimates for both replicates ($C_t < 15$ for telomere and $C_t < 27$ for β-actin). The second quality threshold retained samples for which the between duplicates $C_t$ difference was < 0.6.

a)

| Tissue type | Nb of samples in qPCR analysis | Nb of samples with successful amplification (Ct threshold) for both duplicates | Nb of samples with Ct difference between duplicates < 0.6 |
|---|---|---|---|
| Fins | 33 | 33 | 33 |
| Scales | 33 | 33 | 32 |
| **Number of individuals excluded at this step** | | **0** | **1** |
| **Number of individuals kept for following analyses** | | | **32** |

b)

| River | Collection year | Total individuals in qPCR analysis | Total samples with successful amplification ($C_t$ threshold) for both duplicates | | | Total samples with between duplicates $C_t$ difference < 0.6 | | |
|---|---|---|---|---|---|---|---|---|
| | | | Telomere | β-actin | Both | Telomere | β-actin | Both |
| Acaena | 2018 | 35 | 34 | 35 | 34 | 28 | 34 | 28 |
| | 2003 | 25 | 25 | 24 | 24 | 20 | 18 | 16 |
| Chateau | 2018 | 66 | 64 | 65 | 63 | 53 | 58 | 49 |
| | 2001 | 31 | 26 | 31 | 26 | 22 | 24 | 21 |
| Nord | 2010 | 58 | 57 | 56 | 56 | 42 | 48 | 38 |
| Olsen | 2018 | 25 | 23 | 25 | 23 | 19 | 22 | 18 |
| Orgues | 2018 | 31 | 31 | 31 | 31 | 28 | 30 | 27 |
| Rohan | 2019 | 32 | 32 | 31 | 31 | 23 | 28 | 21 |
| | 2002 | 35 | 35 | 34 | 34 | 22 | 29 | 18 |
| Studer | 2011 | 70 | 70 | 70 | 70 | 57 | 61 | 52 |
| | 2002 | 34 | 34 | 34 | 34 | 22 | 31 | 23 |
| **Total** | | **442** | **431** | **436** | **426** | **336** | **383** | **311** |
| **Number of samples excluded at this step** | | | **16** | | | **115** | | |
| **Number of samples remaining after quality criteria** | | | | | | **311** | | |

### Supplementary Figure S1. Histogram of $C_t$ difference between duplicates for telomere and β-actin.

The vertical grey line indicates the between duplicates $C_t$ difference threshold (0.6). Only samples for which both duplicates showed $C_t$ values lower than 15 (telomere) or 27 (β-actin) are displayed.

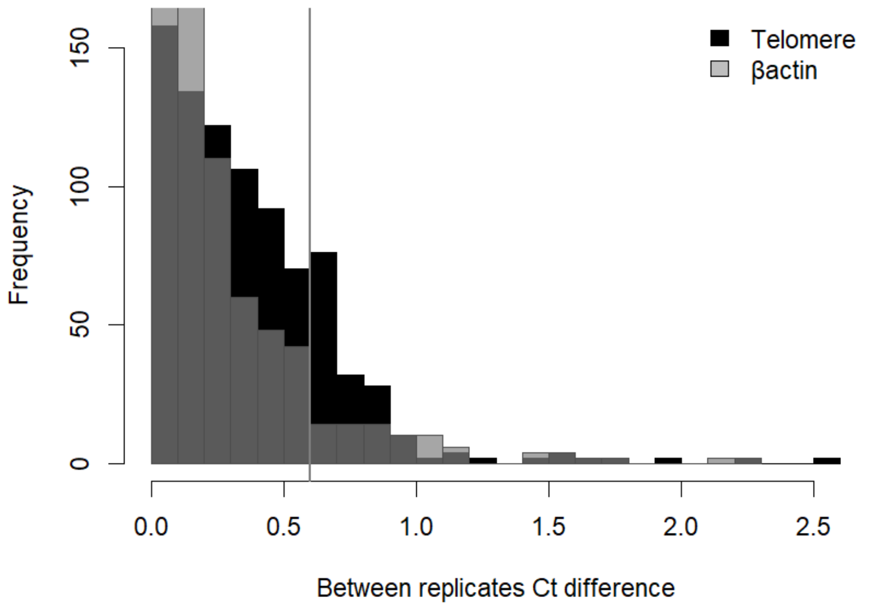

### Supplementary Figure S2. Histogram of all the $C_t$ values measured in Kerguelen samples.

The $C_t$ thresholds of 15 (telomere) and 27 (β-actin) are indicated by the vertical lines.

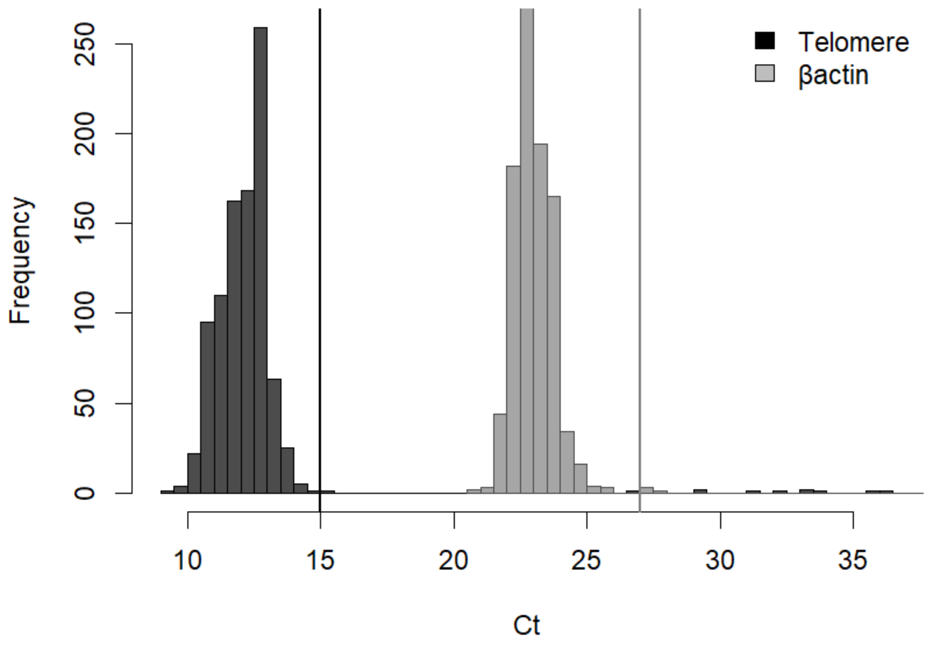

Supplementary Table S4. Results of the GLM model for RTL variation in Kerguelen samples as function of individual age, body size and river of origin.

a) Deviance analysis table

| **Model parameters** | **Degrees of freedom** | **Deviance** | **Residual deviance** | **p-value ($\chi^2$ test)** |
|---|---|---|---|---|
| null | | | 50.759 | |
| river | 6 | 13.808 | 36.951 | $< 2.2\ 10^{-16}$ *** |
| individualAge | 2 | 6.166 | 30.785 | $< 2.2\ 10^{-16}$ *** |
| bodySize | 1 | 0.539 | 30.247 | 0.006 ** |
| river:individualAge | 8 | 10.455 | 19.792 | $< 2.2\ 10^{-16}$ *** |
| river:bodySize | 6 | 0.444 | 19.348 | 0.401 |
| individualAge:bodySize | 2 | 0.335 | 19.013 | 0.096 |
| river:individualAge:bodySize | 7 | 0.777 | 18.235 | 0.145 |

b) Parameter estimates values

| Parameter | Estimate | Std. Error | z value | p-value (Wald z-test) |
|---|---|---|---|---|
| (Intercept) | 0.512 | 0.124 | 4.112 | $5.18\ 10^{-5}$ *** |
| riverChateau | -0.301 | 0.148 | -2.028 | 0.043 * |
| riverNord | -1.472 | 0.330 | -4.457 | $1.21\ 10^{-5}$ *** |
| riverOlsen | -0.605 | 0.190 | -3.178 | 0.002 ** |
| riverOrgues | -0.593 | 0.212 | -2.797 | 0.005 ** |
| riverRohan | -0.302 | 0.135 | -2.233 | 0.026 * |
| riverStuder | -0.277 | 0.156 | -1.776 | 0.077 |
| individualAge2 | -0.412 | 0.136 | -3.035 | 0.003 ** |
| individualAge3 | -0.736 | 0.327 | -2.248 | 0.025 * |
| bodySize | 0.249 | 0.189 | 1.317 | 0.189 |
| riverChateau:individualAge2 | -0.532 | 0.189 | -2.811 | 0.005 ** |
| riverNord:individualAge2 | 0.799 | 0.505 | 1.583 | 0.114 |
| riverOlsen:individualAge2 | NA | NA | NA | NA |
| riverOrgues:individualAge2 | NA | NA | NA | NA |
| riverRohan:individualAge2 | 0.420 | 0.420 | 1.000 | 0.318 |
| riverStuder:individualAge2 | -0.095 | 0.207 | -0.460 | 0.645 |
| riverChateau:individualAge3 | -0.022 | 0.462 | -0.047 | 0.962 |
| riverNord:individualAge3 | 0.589 | 0.559 | 1.053 | 0.293 |
| riverOlsen:individualAge3 | 2.640 | 1.587 | 1.664 | 0.097 |
| riverOrgues:individualAge3 | NA | NA | NA | NA |
| riverRohan:individualAge3 | 0.942 | 0.468 | 2.014 | 0.045 * |
| riverStuder:individualAge3 | NA | NA | NA | NA |
| riverChateau:bodySize | 0.017 | 0.214 | 0.082 | 0.935 |
| riverNord:bodySize | -0.579 | 0.399 | -1.451 | 0.148 |
| riverOlsen:bodySize | -0.299 | 0.277 | -1.079 | 0.281 |
| riverOrgues:bodySize | -0.134 | 0.295 | -0.455 | 0.649 |
| riverRohan:bodySize | -0.332 | 0.220 | -1.511 | 0.132 |
| riverStuder:bodySize | -0.039 | 0.241 | -0.160 | 0.873 |
| individualAge2:bodySize | -0.338 | 0.245 | -1.381 | 0.168 |
| individualAge3:bodySize | -0.300 | 0.212 | -1.413 | 0.159 |
| riverChateau:individualAge2:bodySize | -0.535 | 0.392 | -1.365 | 0.173 |
| riverNord:individualAge2:bodySize | 0.889 | 0.523 | 1.701 | 0.090 |
| riverOlsen:individualAge2:bodySize | NA | NA | NA | NA |
| riverOrgues:individualAge2:bodySize | NA | NA | NA | NA |
| riverRohan:individualAge2:bodySize | 0.567 | 0.537 | 1.057 | 0.291 |
| riverStuder:individualAge2:bodySize | 0.114 | 0.309 | 0.369 | 0.712 |
| riverChateau:individualAge3:bodySize | 0.145 | 0.373 | 0.389 | 0.697 |
| riverNord:individualAge3:bodySize | 1.041 | 0.448 | 2.323 | 0.021 * |
| riverOlsen:individualAge3:bodySize | NA | NA | NA | NA |
| riverOrgues:individualAge3:bodySize | NA | NA | NA | NA |
| riverRohan:individualAge3:bodySize | 0.284 | 0.286 | 0.990 | 0.323 |
| riverStuder:individualAge3:bodySize | NA | NA | NA | NA |
